# Southeast and Northeast facing slopes have the Least Tree Cover in Northern and Southern Tropics, respectively

**DOI:** 10.1101/2025.06.05.658062

**Authors:** D. Maheshwori, S. R. Managave, G. Jathar, S. Davande

## Abstract

Reforestation in the tropics, one of the most deforested regions, can help mitigate climate change and conserve biodiversity. Its effective implementation requires assessing suitability of a given site for tree growth. By analyzing tree cover (TC) in 127 protected areas across the tropics, we show that southeast and northeast-facing slopes are least favorable for tree growth in the Northern and Southern tropics, respectively. In contrast, northwest (north-, northwest, and west-facing) and southwest (south-, southwest, and west-facing) slopes are more suitable for tree growth in the Northern and Southern hemispheres, respectively. This pattern likely results from the combined effects of Pole-Equator and West-East asymmetries in TC. Tropical reforestation programs, especially in hilly areas with intermediate TC, could benefit from considering the influence of slope direction on tree growth and adopting appropriate plantation strategies.

## 1 Introduction

The tropics is one of the most deforested regions (Turubanova et al., 2018; Vancutsem et al., 2021). The forests in the tropics not only constitute a significant sink and storage of carbon (Pan et al., 2011; Lewis et al., 2019) but also have potential for sequestering additional carbon through regeneration (Williams et al., 2024). Increase in tree cover (TC) in tropics through afforestation and reforestation can mitigate the effects of climate change (Bastin et al., 2019). Additionally, reforestation can help in conserving biodiversity, preventing soil erosion and providing ecosystem services (Locatelli et al., 2015; Chazdon and Brancalion, 2019). Numerous national and international initiatives have aimed at increasing tropical forest restoration e.g. the 2030 Agenda for Sustainable Development (2030 Agenda), UN Decade on Ecosystem Restoration (2021–2030) and the UN Framework Convention on Climate Change (UNFCCC).

Identification of the factors that facilitate or hamper establishment of TC is important for the successful implementation of programs that aim to increase TC. The mean annual precipitation (Sankaran et al., 2005; Hirota et al., 2011) and its variability (Holmgren et al., 2013; Xu et al., 2018) are known to control spatial distribution of TC in the tropics. The local topographical drivers can put additional spatially heterogeneous constraints on tree growth at the landscape-scale (Geroy et al., 2011; Gutiérrez-Jurado et al., 2013; Zhou et al., 2013; Werner & Homeier 2015).

The direction a given slope faces (also called aspect) and its slope angle are important topographical drivers of vegetation at the landscape-scale (Singh, 2018; Smith and Bookhagen, 2021). Equator-facing slopes receive higher solar insolation than Pole-facing slopes resulting in it having relatively warmer, drier microclimate and soil with lower moisture content than Pole-facing slopes (Geroy et al., 2011; Gutiérrez-Jurado et al., 2013; Zhou et al., 2013). As a result, Equator-facing slopes have a lower vegetation cover than Pole-facing slopes, and Pole-Equator asymmetry (Holland and Steyn, 1975; Badano et al., 2005; Singh, 2018; Smith and Bookhagen, 2021). In addition, it has been also shown that West-facing slopes have higher vegetation cover than the East-facing slopes (Smith and Bookhagen, 2021), the West-East asymmetry. Both the asymmetries are commonly observed at mid-latitude regions (e.g. Smith and Bookhagen, 2021). However, their influence on tree growth in tropics is not fully recognized.

Selection of an appropriate area is important for successful reforestation (Sacco et al., 2020). The demonstration of the Pole-Equator and West-East asymmetries (Smith and Bookhagen, 2021; Maheshwori et al., 2025) highlights the existence of heterogeneous conditions for tree growth on different aspects at the landscape scale. However, how and where these two modes influence TC on different aspects in the global tropics is still incomplete. Here, using TC data in 127 protected areas (PAs) distributed throughout the tropics, we investigate (i) conduciveness of different slope directions for tree growth and (ii) the regions where slope aspect exerts a significant control over tree growth.

## 2 Data and methodology

### 2.1 Data sets

To examine the influence of slope direction on TC in tropical regions, we used the Hansen Global Forest Change v1.9 (2000–2021) dataset (Hansen et al., 2013) which provides percent tree cover in 30 m x 30 m grid cells. The Shuttle Radar Topography Mission (SRTM) Digital Elevation Model (DEM) with 30 m resolution (NASA JPL, 2013) was used for topographical analysis, including elevation, slope, and aspect.

### 2.2 Methodology

To understand the effect of slope-aspect on TC, we analysed TC in 361 PAs distributed throughout the tropical hilly areas. The PAs, selected from *The Protected Planet: World Database on Protected Areas (WDPA)* (UNEP-WCMC & IUCN, 2025), were selected to minimize the effect of anthropogenic activities. For analytical efficiency, geographically proximate PAs were considered together which reduced the number of PAs to 127. The data for selected PAs in India (N = 25) were from Maheshwori et al., (2025). To assess the effect of absolute TC on the aspect-related heterogeneity in TC, PAs with a wide range of median TC (ranging from 0 to 100%) were selected.

Topographic parameters viz. slope and aspect were computed from SRTM DEM using QGIS (QGIS Development Team, 2023). The TC data, available in TIFF format, were converted to a gridded data format in QGIS. To align the TC data with the corresponding DEM coordinates, the nearest-neighbor re-gridding method was used. We examined mean and median TC variations on eight aspect classes i.e. North (N), Northeast (NE), East (E), Southeast (SE), South (S), Southwest (SW), West (W), and Northwest (NW). The TC on four slope angle (0°–10°, 10°–30°, 30°–50°, and 50°–90°), and five elevation (0-200 m, 200–500 m, 500–900 m, 900–1500 m, and 1500–4000 m) bins were also analysed.

### 2.3 Statistical treatment

The TC of entire PAs were analysed i.e. 100% sampling. The TC distribution on different slope direction in individual PAs had different skewness (range: 0.07 to 6.27; mean 1.35 ± 1.28, 1-sigma). A non-parametric test (two-sample Kolmogorov-Smirnov test) showed that the distributions of TC on different aspects in 123 PAs (out of 127) were statistically different (p < 0.05). In four PAs (from Australia where the median TC were 0%) the distributions were statistically similar (p > 0.05). We used median TC while comparing TC on different aspects. Student’s t-test (two-tailed) was performed while assessing significance of the difference between two means.

## 3 Results and Discussion

### 3.1 Pole-Equator and West-East asymmetries in TC

The TC distribution in the PAs showed the expected increase in the Pole-Equator and West-East asymmetries with latitude (Fig. 2) and slope (Suppl. Fig. S1, S2, S3). The TC on the Pole-facing slopes were similar to that on West-facing slopes (−0.28 ± 2.25). However, the TC on Equator-facing slopes were higher than West-facing slopes (5.37 ± 6.21) implying the Pole-Equator asymmetry is generated by selectively lowering TC on Equator-facing slopes.

**Figure 1.**
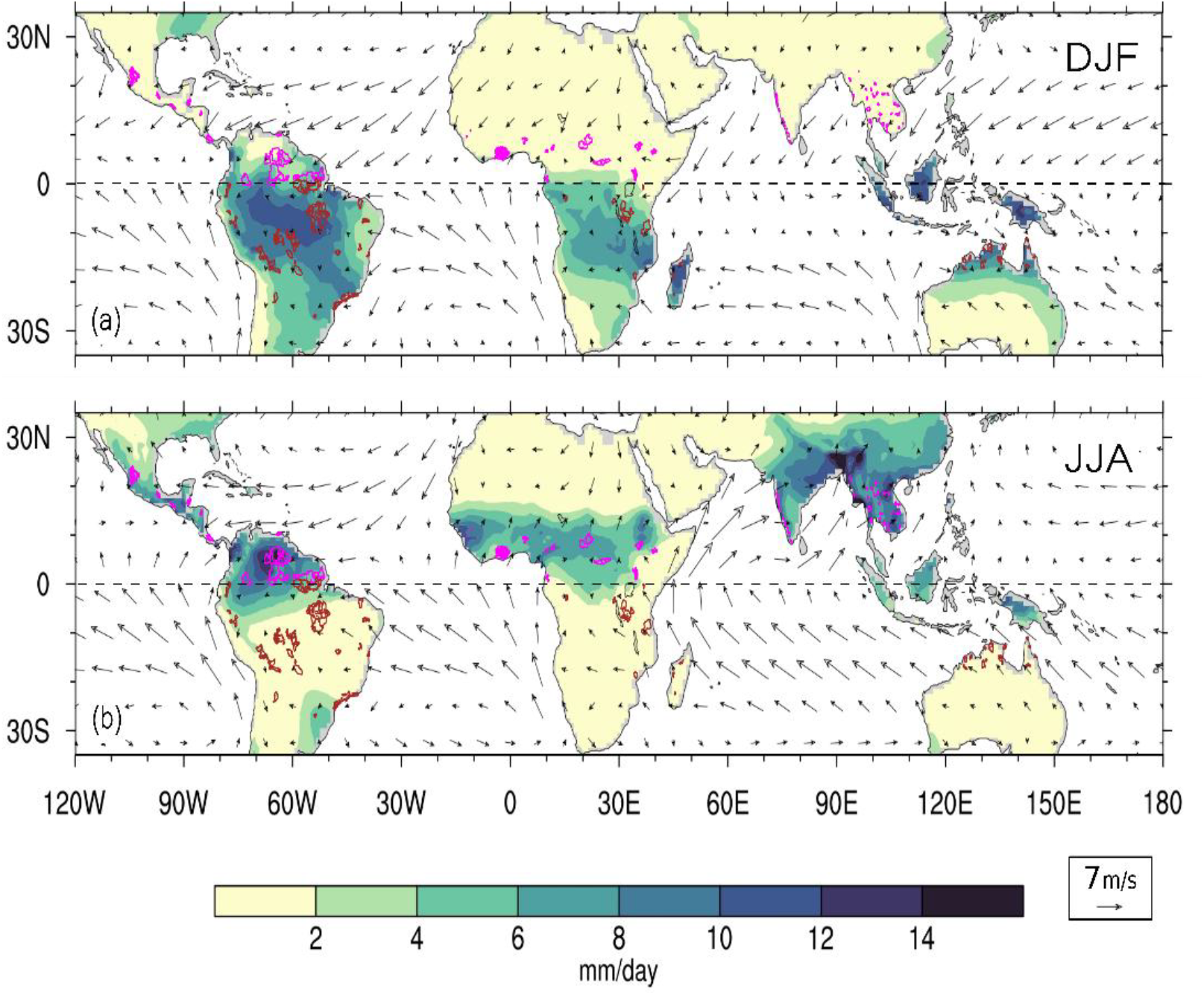
Protected areas (outlines marked in magenta and red) whose tree cover was analysed in this study. Precipitation (Global Precipitation Climatology Project Daily Precipitation (Huffman et al., 2001; Adler et al., 2017)) and wind climatology (NCEP/NCAR reanalysis surface wind speed) (Kalnay et al., 1996) during December to February (DJF) (a) and June to August (JJA) (b).

**Figure 2.**
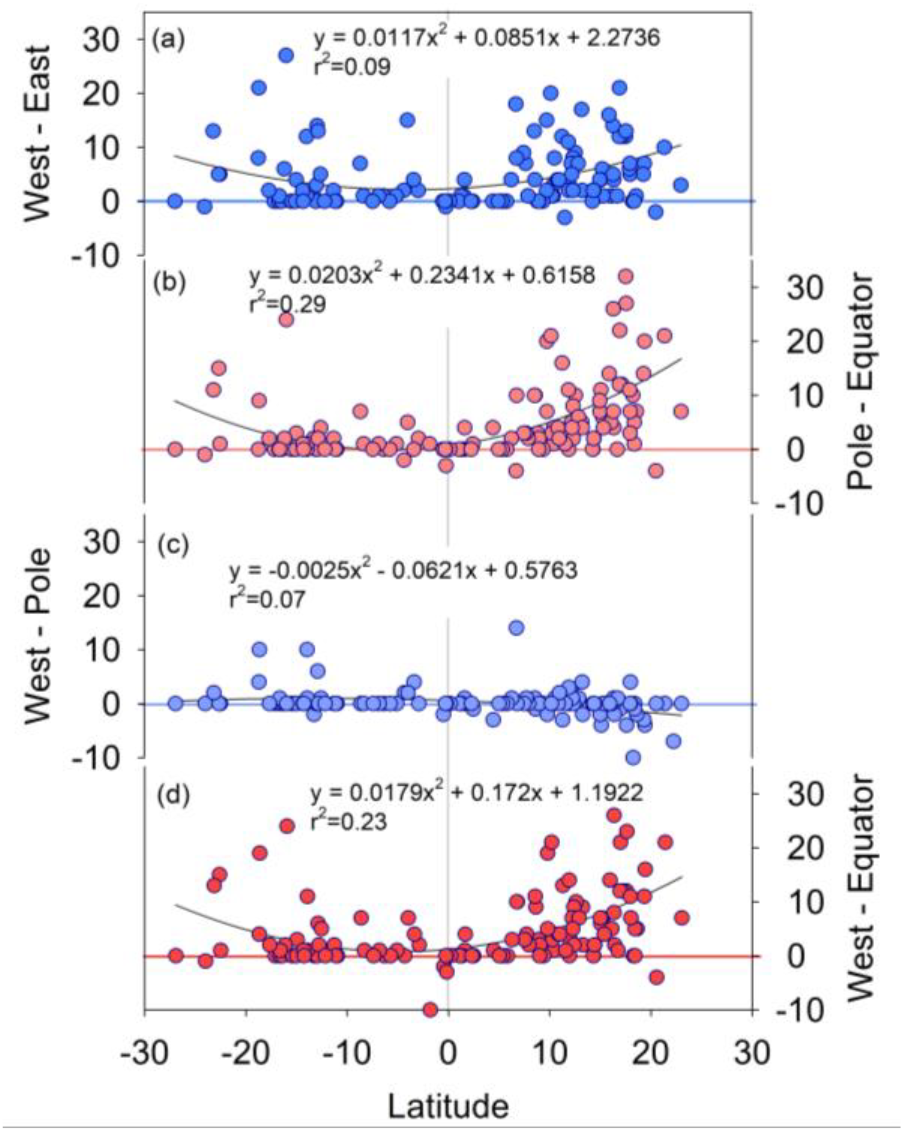
Latitudinal variation in the difference between median TC on slopes facing West and East (a), Pole and Equator (b), West and Pole (c), and West and Equator (d).

The Pole-Equator asymmetry is attributed to the asymmetrically higher solar insolation received by Equator-facing slopes than Pole-facing slopes (Smith and Bookhagen, 2021) which leads to the former having warmer, drier microclimate with lower soil moisture (Måren et al., 2015; Burnett et al., 2008). The duration the Sun spends over Equator-facing slopes is higher than over Pole-facing slopes; this asymmetry increases poleward from the equator (Maheshwori et al., 2025). While this asymmetry in the insolation (i.e. modelled clear-sky solar radiation at a surface) explains the asymmetric heating of the Pole- and Equator-facing slopes (Smith and Bookhagen, 2021), the plausible role played by rain and clouds in further enhancing the asymmetry in the radiation received on both faces is under-appreciated (Fig. 1 and Fig. 3). The Inter Tropical Convergence Zone (ITCZ) and associated rain-bearing clouds reside in the Northern Hemisphere (NH) during boreal summer (Fig. 1). During this time, the Pole- and Equator-facing slopes in the NH receive rains and lesser solar radiation due to the clouds (as manifested in lower ongoing longwave radiation (Waliser et al., 1993). The Southern Hemisphere (SH) is nearly cloud free and dry (Fig. 1 and 3) with its Equator-facing slopes getting heated preferentially during boreal summer. The preferential heating of the Equator-facing slopes in NH starts since the ITCZ starts moving southwards (e.g. ∼October at 20° N) and continues till the return of the ITCZ (e.g. ∼June at 20° N). The opposite scenario occurs during Austral summer, which leads to heating of equator facing slopes in NH (Fig. 3). Thus the combined effect of the Sun spending longer duration over Equator-facing slopes and a concurrent dry period leads Equator-facing slopes to receive higher solar radiation which results in their lower TC.

**Figure 3.**
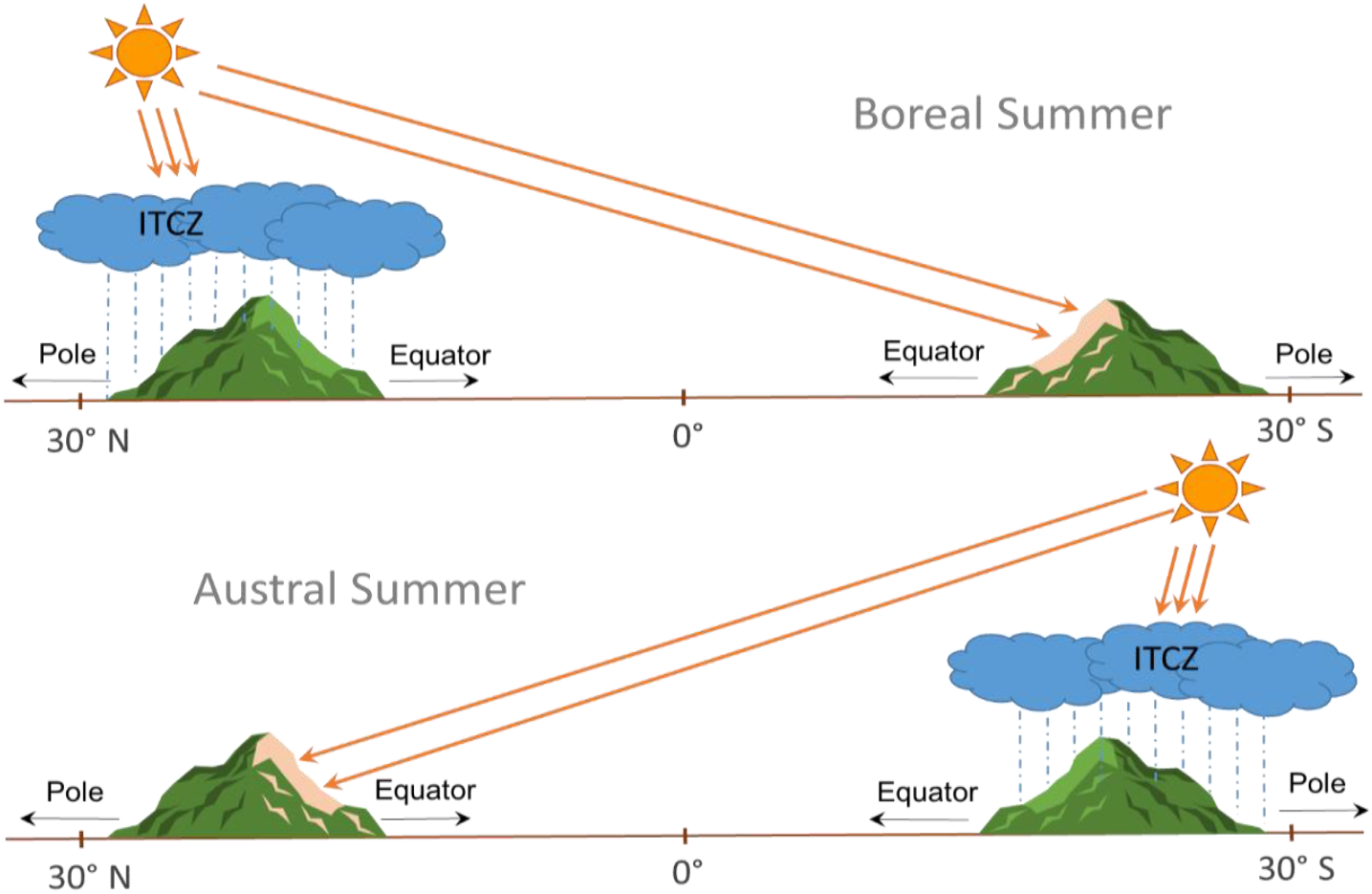
A schematic diagram showing the role of solar inclination and ITCZ position in establishing Pole-Equator asymmetry in the solar radiation received.

As the West and East facing slopes receive the similar intensity of insolation (i.e. clear-sky solar radiation) (Smith and Bookhagen, 2021), it cannot explain the observed West-East symmetry in TC. Orography can induce precipitation asymmetry on West- and East-facing slopes. For example, in Asia, the monsoonal winds flow from the Southwest direction resulting in a windward West-facing region receiving higher precipitation than leeward East-facing region (Maheshwori et al., 2025). The other factors such as persistence of morning dew periods, timing of the maximum cloud cover and the ambient air temperature during peak sunlight hours could explain the West-East asymmetry in vegetation (Smith and Bookhagen, 2021) and the observed TC.

### 3.2 Variation of absolute TC on Pole-, Equator, West- and East-facing slopes

The slope-wise variations in the Pole-Equator and West-East asymmetries only indicate the difference in the TCs with slope. They do not reveal how actual TC varies with slope on these aspects which is more important for assessing variation in the conduciveness for tree growth with slopes on these aspects. The TC on West- and Pole-facing slopes was higher on the steeper slopes than on 0°-10°; the TC increased with slope from 0°-10° to 30°-50° and slightly decreased thereafter (Fig. 4 a and c). In contrast, the TC on East and Equator-facing slopes increased only slightly from 0°-10° to 30°-50°; and thereafter it decreased with slope (Fig. 4 b and d). The TC on slopes from 30° to 90° were lower than that on 0°-10° on East- and Equator-facing slopes (Fig. 4 b and d). Thus the increase in the Pole-Equator and West-East asymmetries in TC with slope is due to selective lowering of TC on Equator- and East-facing slopes.

**Figure 4.**
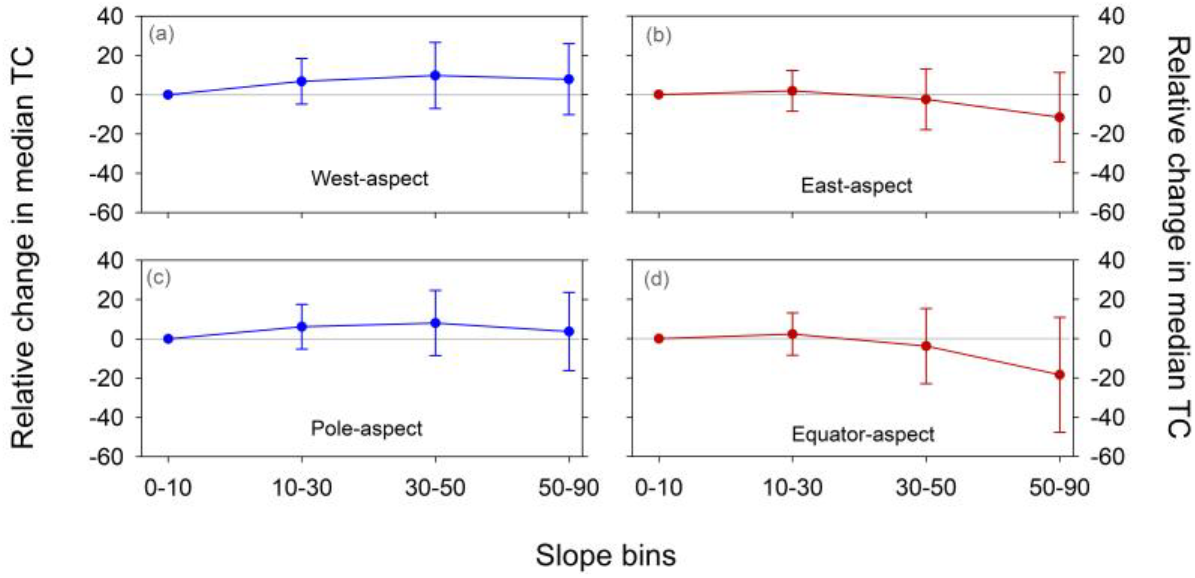
Change in the median TC with slope relative to median TC in 0°-10° bin on West-(a), East-(b), Pole-(c) and Equator-(d) facing slopes. The points represent mean and 1-sigma standard deviations of relative changes in the median TC with slope in PAs. The changes in TC (relative to TC in 0°-10° bin) on West- and East-facing slopes, (and on Pole- and Equator-facing slopes) in their respective slope bins were statistically different (P < 0.006).

### 3.3 TC on various aspects

The slope direction which showed the highest and lowest TC varied in the NH and SH (Fig. 5 and Suppl. Fig. S4). The lowest TC was observed on SE- and NE-facing slopes in the NH and SH, respectively; the highest TC was on NW-facing slopes in the NH and, W-facing slopes in the SH (Fig. 5). Similar patterns were also observed in individual regions (Fig. 6, Suppl. Fig. S5). We attribute this to the combined effect of the Pole-Equator and West-East modes that influence the TC at the landscape-scale. In the NH, the decreasing trend in TC was from North-to South-facing slopes and from West-to East-facing slopes (Fig. 2 and Maheshwori et al., 2025). The polarity of only the former reverses in SH i.e. decreasing trend in TC from South-to North-facing slopes. As a result, the SE-facing slopes in the NH and NE-facing slopes in the SH have the disadvantage of both modes of TC variability and have the lowest TC. On the similar lines, NW and SW facing slopes in the NH and SH, respectively, have advantage of both modes and have the higher TC.

**Figure 5.**
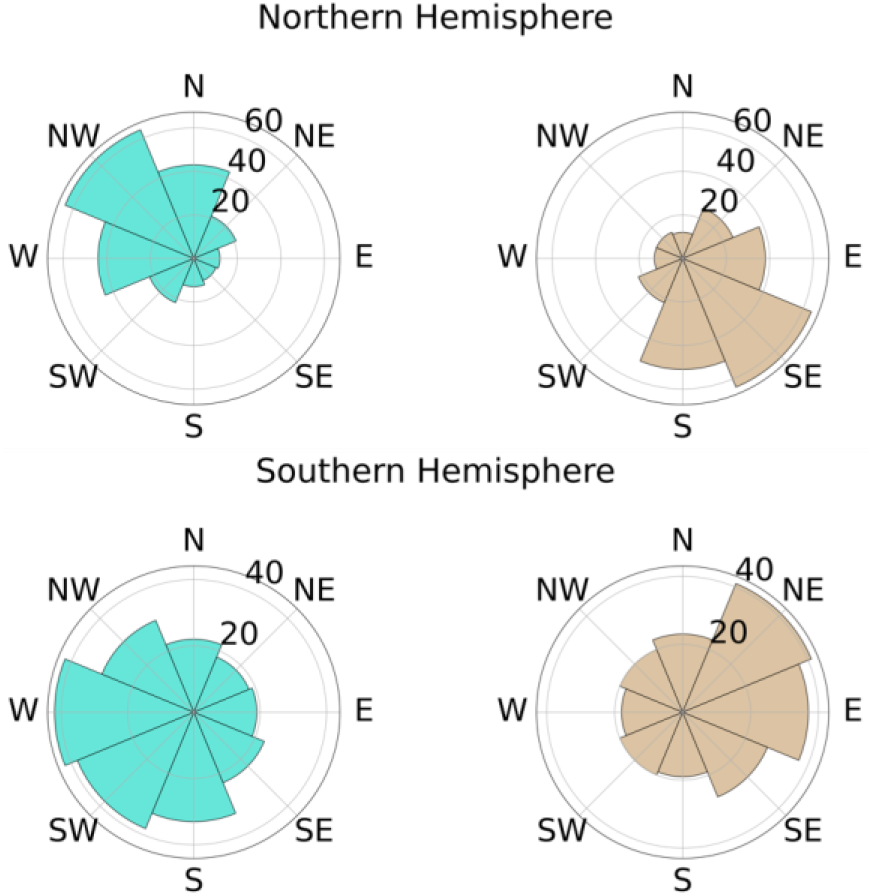
The number of PAs that showed the highest (blue) and lowest (brown) median tree cover in different aspect categories in the Northern (top panels) and Southern (bottom panels) hemispheres.

**Figure 6.**
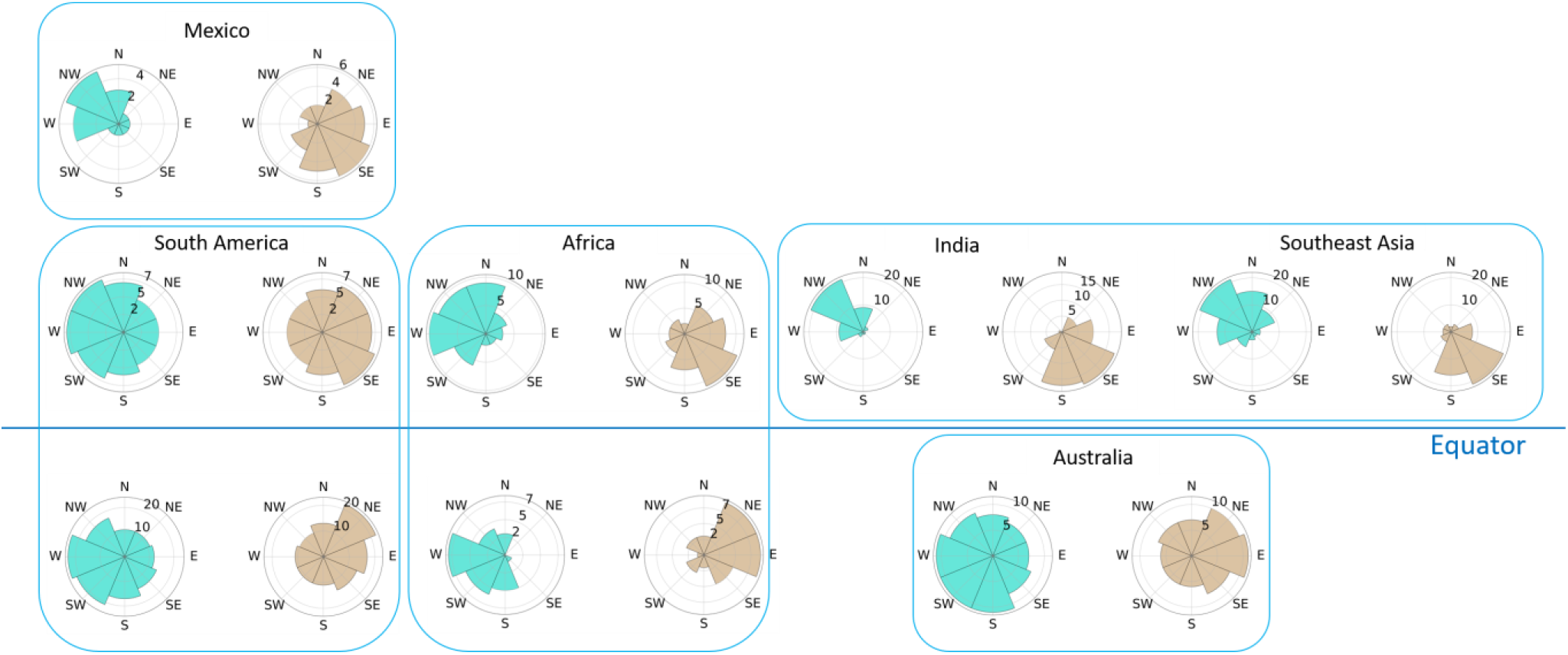
The number of PAs that showed the highest (blue) and lowest (brown) median tree cover in different aspect categories in various regions.

The maximum difference in the TC was observed between the slopes facing NW-(N-, NW- and W-facing slopes) and SE-quadrants (S-, SE- and E-facing slopes) in the NH; most of the PAs showed the highest difference between NW- and SE-facing slopes (Fig. 7a). The opposite was observed in the SH: the maximum difference in TC was between the slopes facing SW-(W-, S- and SW-facing slopes) and NE-quadrants (N-, NE- and E-facing slopes) (Fig. 7b). The similar pattern was observed at the regional scale (Suppl. Fig. S6). This is in contrast to previously reported pattern that suggested the greatest aspect differences in vegetation cover and/or biomass generally occurs between SW-facing versus NE-facing hillslopes in the NH and between NW facing versus SE-facing hillslopes in the SH (Boyko, 1947; Perring, 1959; Holland and Steyn, 1975; Radcliffe and Lefevre, 1981; Bale et al., 1998; Desta et al., 2004; Pelletier and Swetnam, 2017).

**Figure 7.**
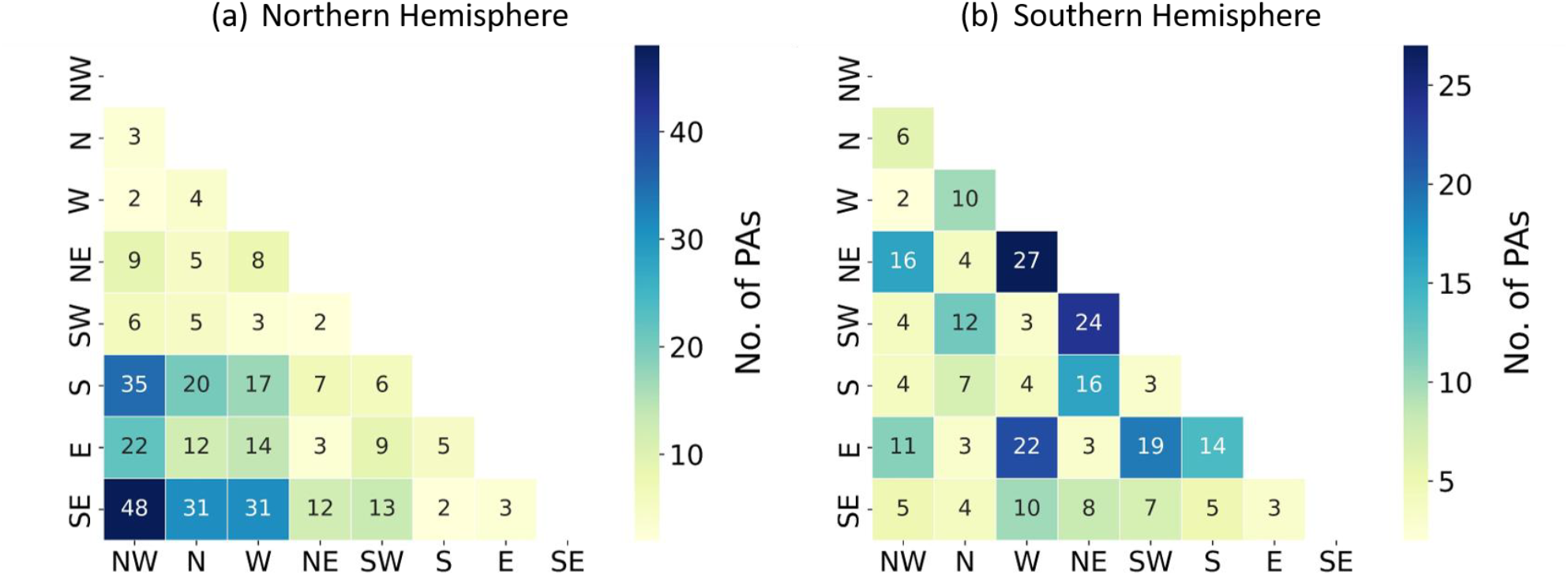
The number of PAs in the Northern (a) and Southern Hemispheres (b) that showed the highest difference in the median TC between various combinations of aspects. While estimating this, the PAs that showed the same median TC in all aspects (N = 23) were ignored.

### 3.4 Relationship of absolute TC with aspect-related heterogeneity in TC within PAs

In addition to the slope, the magnitude of maximum difference in TC on different aspects in the PAs was also influenced by the mean TC in the PAs (Fig. 8). When the conditions were too conducive (i.e. mean TC ∼90 to 100%) or too unfavorable (i.e. mean TC <10 %) for tree growth the aspect-related heterogeneity in the decreased (Fig. 8). The regions with the intermediate TC (10 to 80%) exhibited the larger differences (Fig. 8).

**Figure 8.**
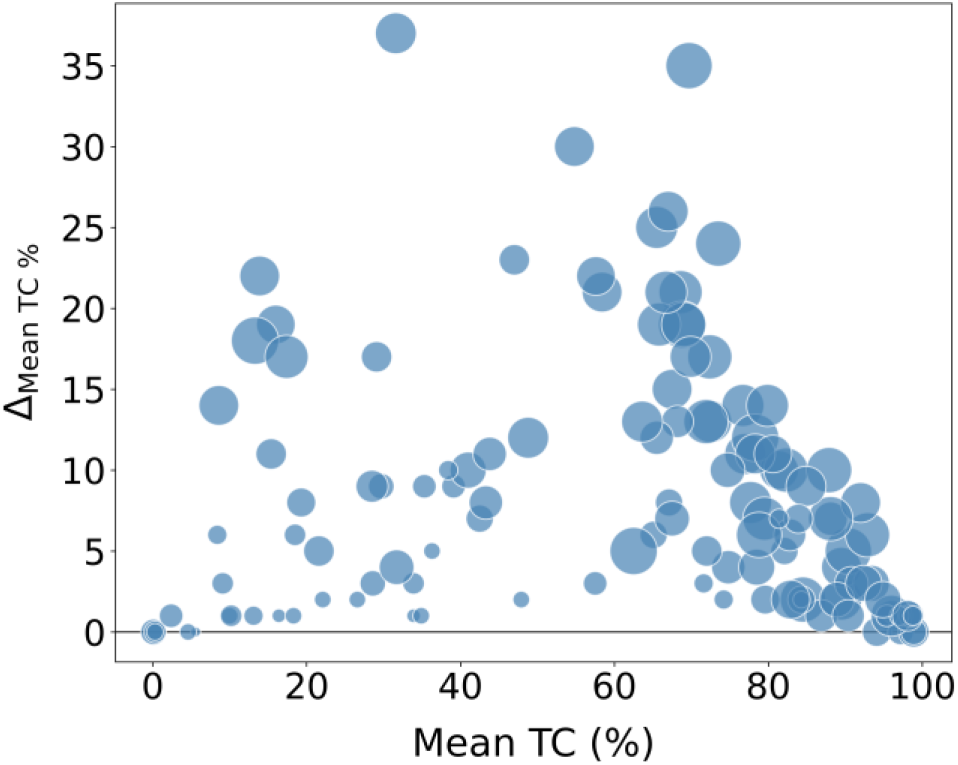
The effect of mean TC in the PAs on the maximum aspect-related difference in mean TC in PAs. The bubble size is proportional to the mean slope in the PA.

## 4 Conclusions

Numerous reforestation programs are being carried out in tropics owing to various drivers such as agroforestry, sustainable forest management, protection of existing forests, park establishment, government sponsored projects (Nagendra and Southworth, 2010). Even though, the structure and composition of TC are known to vary significantly across small spatial scales (Werner & Homeier 2015; Sullivan et al. 2017; Jucker et al., 2018), how the slope-aspect influence TC at the landscape scale is still evolving. We show variations in conduciveness for tree growth on various aspects. Reforestation programs in the tropical hilly regions having intermediate TC (10 to 80%) should consider the following.

1. It is important to recognize that certain aspects (the SE-aspects in the NH and NE-aspects in the SH) have relatively less favorable conditions for tree growth; their TC is not ‘secondary/degraded’ while that on other (N-, NW- and W-aspects in the NH, and W- and SW-aspects in SH) is ‘primary/pristine’.
2. The reforestation that aim to create higher TC on the aspects that are less favorable for tree growth (e.g. that aim to mimic TC of NW-aspect on SE-aspect in the NH) are likely to be expensive and less effective. Such efforts are less likely to succeed and might lead to negative impact on local hydrology as sustaining trees cover through reforestation where water demand is higher than that can be met by the rains alone can lead to local water scarcity (Ricciardi et al., 2022).
3. The ‘targets’ for increasing TC through reforestation/afforestation are likely to be different on different aspects. We recommend considering slope aspect as one of the parameters while estimating ecological restoration potential of an area, formulating reforestation strategy and assessing success of the ongoing reforestation programs.

## Acknowledgments

Help extended by Rahul Tak with data handling and Pratiksha Baruah with wind and precipitation climatology plots are appreciated.

## Author contributions

Conceptualization: Shreyas Managave

Methodology: Shreyas Managave, Devi Maheshwori, Girish Jathar, Sham Davande

Formal Analysis: Devi Maheshwori, Shreyas Managave, Girish Jathar, Sham Davande

Investigation: Devi Maheshwori, Shreyas Managave, Girish Jathar, Sham Davande

Resources: Shreyas Managave

Writing—Original Draft: Shreyas Managave, Devi Maheshwori

Writing—Review and Editing: Shreyas Managave, Devi Maheshwori, Girish Jathar, Sham Davande

Supervision: Shreyas Managave

Funding Acquisition: Shreyas Managave

## Competing interests

The authors declare no competing interest

## Funding

This work was supported by Anusandhan National Research Foundation (ANRF), India (Grant: CRG/2022/005854).

## Data Availability

Data will be made available upon request to the corresponding author.

## Notes

### Competing Interest Statement

The authors have declared no competing interest.

