## Supplementary Figures for "Southeast and Northeast facing slopes have the Least Tree Cover in Northern and Southern Tropics, respectively"

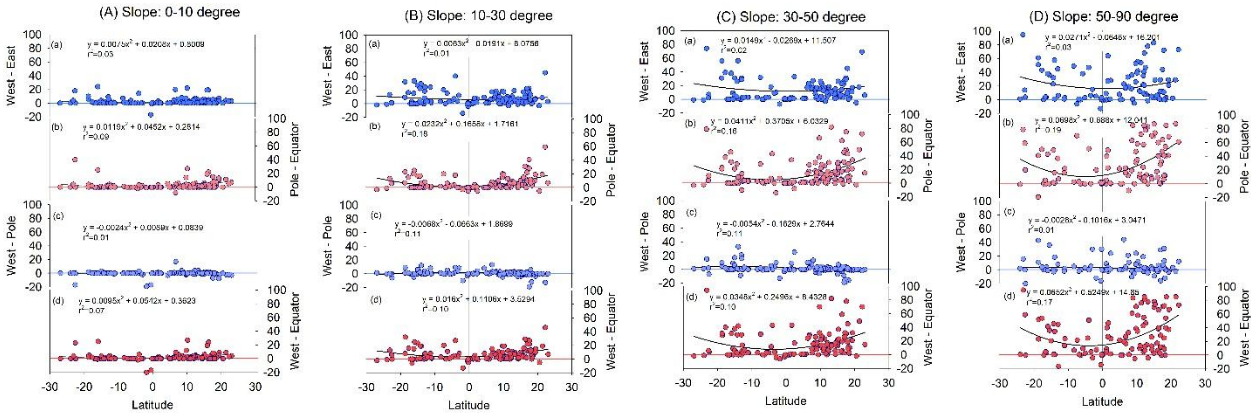


*Supplementary Figure S1. The effect of slope (A to D) on the latitudinal variation in the difference between median TC on slopes facing West and East (a), Pole and Equator (b), West and Pole (c), and West and Equator (d).*

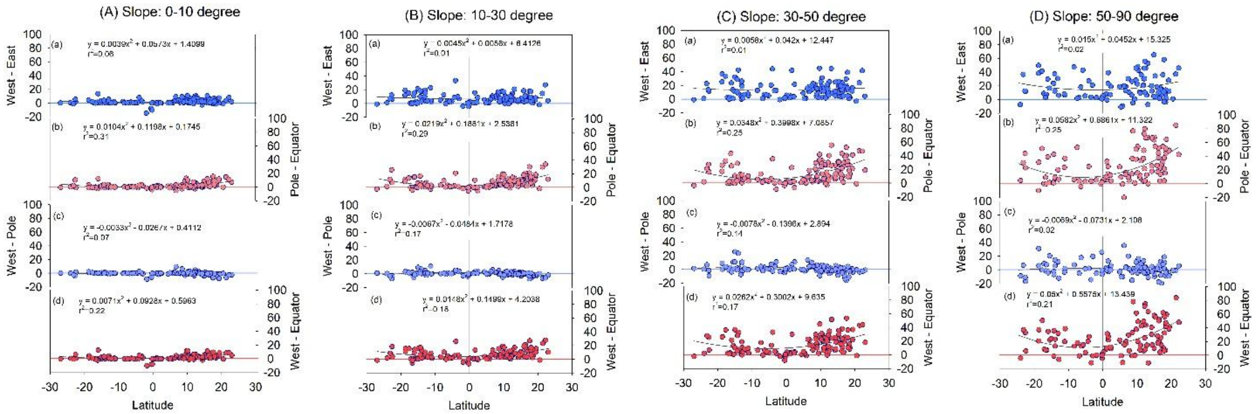


*Supplementary Figure S2. The same as Suppl. Fig. S1 but for mean TC.*

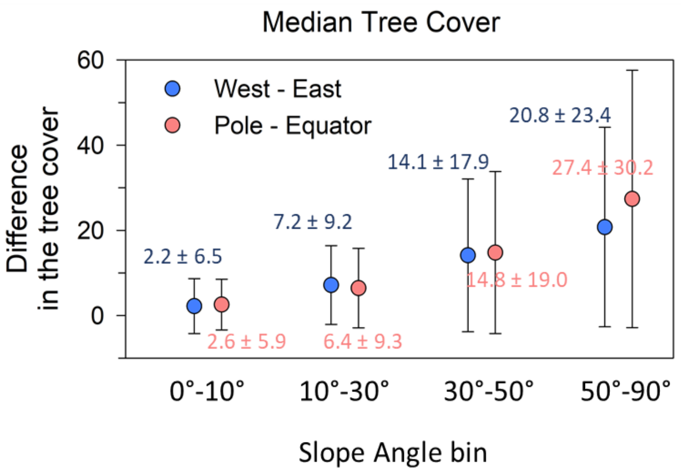


*Supplementary Figure S3. Slope-wise variations in the difference between median TC on West- and East-facing (blue circles), and between Pole-Equator facing (pink circles) slopes. The error bar indicates 1-sigma standard deviations. The differences of each (West-East or Pole-Equator) with slope were significant at P<0.02.*

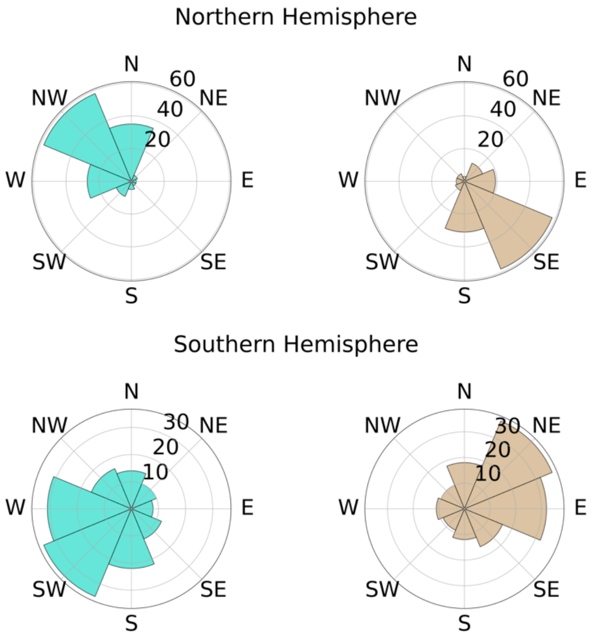


*Supplementary Figure S4. The same as Figure 5 in the main text but for the mean TC. The numbers of PAs showing the highest (blue) and lowest (brown) mean tree cover in different aspect categories in the northern (top panels) and southern (bottom panels) hemispheres.*

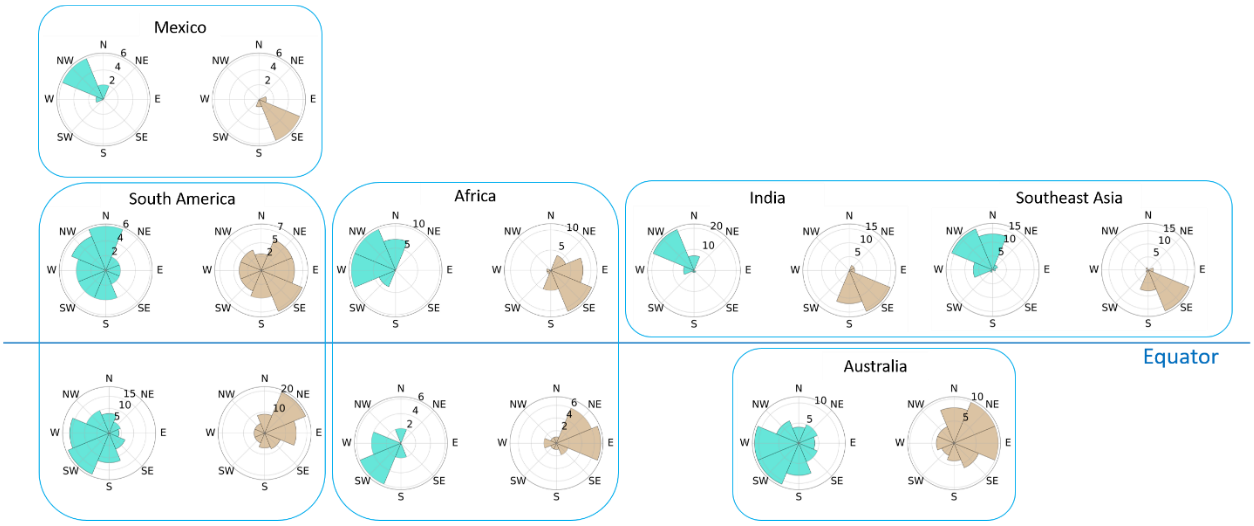


*Supplementary Figure S5. The same as Figure 6 in the main text but for the mean TC. Percent PAs showing the highest (blue) and lowest (brown) mean tree cover in different aspect categories in various regions.*

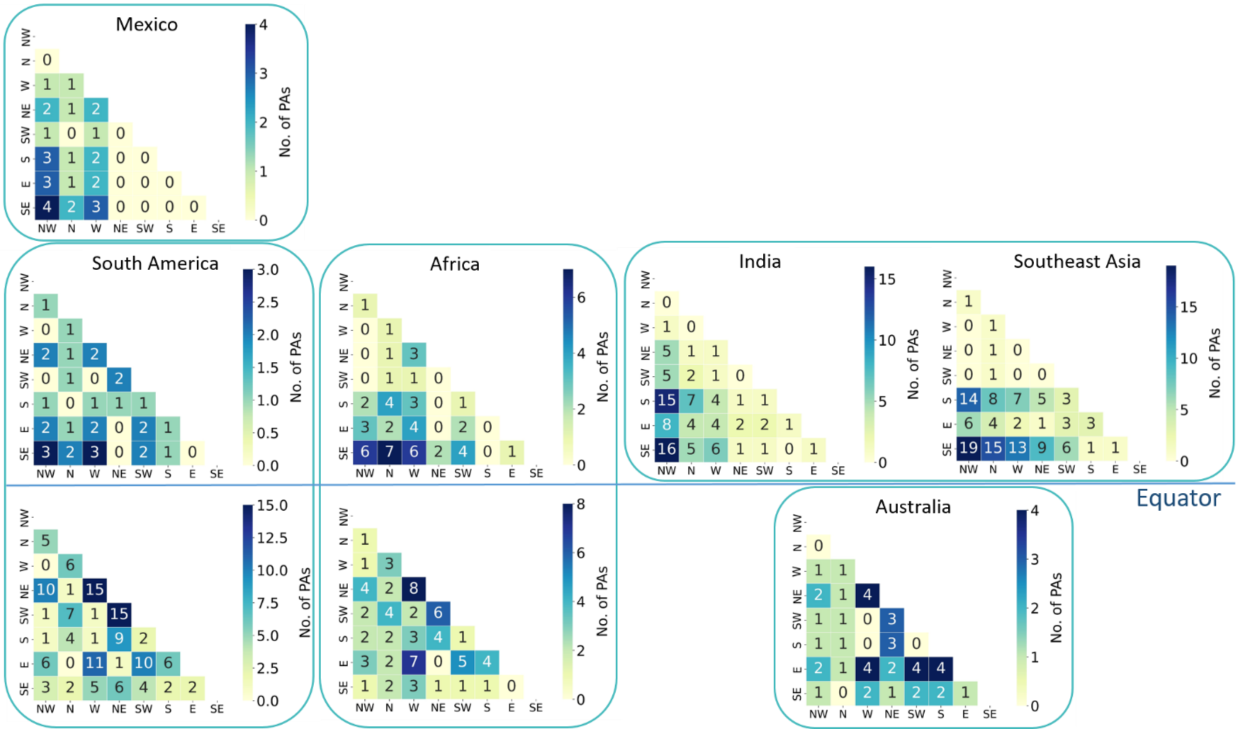


*Supplementary Figure S6: Region-wise number of PAs that showed the highest difference in the median TC between various combinations of aspects. While estimating this, the PAs that showed the same median TC in all aspects were ignored.*
